# Proteomic Sensors for Quantitative, Multiplexed and Spatial Monitoring of Kinase Signaling

**DOI:** 10.1101/2024.12.16.628391

**Authors:** William J Comstock, Marcos VAS Navarro, Deanna V Maybee, Yiseo Rho, Mateusz Wagner, Yingzheng Wang, Marcus B Smolka

## Abstract

Understanding kinase action requires precise quantitative measurements of their activity *in vivo*. In addition, the ability to capture spatial information of kinase activity is crucial to deconvolute complex signaling networks, interrogate multifaceted kinase actions, and assess drug effects or genetic perturbations. Here we developed a proteomic kinase activity sensor platform (ProKAS) for the analysis of kinase signaling using mass spectrometry. ProKAS is based on a tandem array of peptide sensors with amino acid barcodes that allow multiplexed analysis for spatial, kinetic, and screening applications. We engineered a ProKAS module to simultaneously monitor the activities of the DNA damage response kinases ATR, ATM, and CHK1 in response to genotoxic drugs, while also uncovering differences between these signaling responses in the nucleus, cytosol, and replication factories. Furthermore, we developed an *in silico* approach for the rational design of specific substrate peptides expandable to other kinases. Overall, ProKAS is a novel versatile system for systematically and spatially probing kinase action in cells.

## Introduction

Most cellular processes are controlled by the coordinated action of protein kinases. However, the multifaceted actions of over 500 kinases in human cells pose major challenges for the study of kinase function and the deconvolution of signaling networks in cells. A range of techniques are available for investigating *in vivo* kinase activity, although each technology suffers from important intrinsic limitations. Mass spectrometry-based phosphoproteomic technologies are capable of mapping and quantitatively analyzing tens of thousands of phosphorylation events in cells, however, they lack the ability to probe kinase action with spatial resolution, a key variable in the study of kinase biology ^1,2^. Phosphoproteomic approaches are also not easily parallelizable, making them less amenable to high throughput analyses. Moreover, phosphoproteomic data do not inform on direct kinase-substrate relationships unless combined with computational motif analyses, whose assignments are often ambiguous and biased towards well-characterized kinases ^3^. Phosphorylation site-specific antibodies have been traditionally used to image the location of a phosphorylation event in cells using microscopy ^4,5^. Unfortunately, antibodies displaying high specificity and low background for a given phosphorylation site are difficult to generate and not widely available. Therefore, experiments based on phospho-specific antibodies commonly suffer from low signal to noise ratio, limited dynamic range, and tend to yield semi-quantitative results that require extensive and complex image analyses and are not compatible with high throughput analytical purposes.

Genetically encoded fluorescent biosensors offer a solution for the direct visualization of kinase activity toward specific target peptides in living cells ^6,7^. Fluorescent kinase sensors, typically designed with a pair of fluorescent proteins for Förster Resonance Energy Transfer (FRET) or a conformationally changing circularly permuted fluorescent protein (cpFP), integrate a kinase-specific peptide substrate with a phospho-motif binding domain ^7–9^. Upon phosphorylation by the kinase of interest, the recognition of the phospho-peptide by the binding domain induces a conformational change, triggering changes in energy transfer in FRET-based sensors or fluorescence in cpFP-based sensors ^9^. Alternatively, the phosphorylation of the peptide substrate sequence can cause translocation or degradation of the fluorophore ^7^. Cell-penetrating fluorescent (FLIM) probes for Abl and Src-family kinases have also been designed and implemented without the need for FRET^10^. Despite successful application of fluorescent kinase sensors in a range of biological contexts, these systems have limitations, both in terms of general applicability as well as technical implementation. The specificity required for recognition by the phospho-motif domain limits the range of kinases that can be monitored, and the biosensor design process is labor-intensive, often requiring extensive optimization ^11^. Moreover, excessive binding affinity between the phospho-motif and the interaction domain can lead to sensor saturation, restricting dynamic range and potentially impeding dephosphorylation by phosphatases, as observed with a designed ATM sensor, which showed reduced efficiency in monitoring phosphorylation decay following kinase inhibition due to excessive binding strength ^12^. Additionally, multiplexing these sensors is constrained by fluorescence signal overlap, limiting the ability to track multiple kinase activities concurrently ^11^.

Damage to DNA or stress during DNA replication trigger the activation of phosphatidylinositol 3′ kinase (PI3K)□related kinases (PIKKs) ATR, ATM and DNA-PKcs ^13–15^. Once activated, PIKKs orchestrate elaborate signaling responses that regulate a range of cellular processes such as DNA repair, DNA replication, cell cycle, transcription, and apoptosis. The downstream kinases CHK1 and CHK2 are activated by ATR and ATM, respectively, and mediate key aspects of the overall signaling response, including replication fork stability and cell cycle progression ^16,17^. Despite extensive studies on PIKKs and downstream signaling responses and the mapping of the signaling network controlled by these kinases, our understanding of the spatial organization of kinase signaling within distinct subnuclear domains and cellular compartments remains elusive. There is currently a need for quantitative tools capable of rigorously and systematically monitoring locations and kinetics of DNA damage signaling in a lesion- and cell type-dependent manner and with high dynamic range and throughput.

Here we developed the Proteomic Kinase Activity Sensor (ProKAS) platform that leverages MS for the multiplexed, spatially resolved, and quantitative monitoring of kinase activity in living cells. ProKAS is based on a tandem array of peptide sensors that allows simultaneous tracking of multiple kinases within a single polypeptide module. The introduction of amino acid barcodes into these peptide substrates enables the multiplexed monitoring of kinase activities across different cellular locations or under varying experimental conditions. The multiplexing capabilities make ProKAS compatible with high throughput analyses and screening purposes. We engineered a ProKAS module specifically designed to sense the activity of DNA damage response kinases ATR, ATM, and CHK1. Our results demonstrate the ability of the ProKAS sensor to capture kinase activity with high specificity and spatio-temporal resolution, uncovering kinetics of kinase signaling during DNA damage responses. Additionally, we developed a *de novo* approach for the rational design of substrate peptides, which is expected to be broadly applicable to most kinases within the human kinome. Overall, ProKAS offers a versatile platform for probing entire kinase signaling networks, opening new avenues for investigating kinase action in cells.

## Results

### ProKAS biosensor design

We designed the Proteomic Kinase Activity Sensor (ProKAS) platform to allow for quantitative, multiplexable and spatially resolved monitoring of kinase activity in cells using mass spectrometry (MS) (Figure 1). The core of ProKAS lies on a Multiplexed Kinase Sensor (MKS) module, a tandem array of 10-15 amino acid peptide sensors representing preferred substrate motifs for selected kinases of interest (KOIs) (Figures 1A-B). Each sensor has a serine or threonine at its center and three residues at the edge for barcoding purposes (Figure 1B). The ProKAS polypeptide also features an N-terminal enhanced Green Fluorescent Protein (eGFP) for imaging and a Targeting Element (TE) that directs the sensor to a specific subcellular location (e.g. nuclear localization signal (NLS), nuclear export signal (NES), or a protein of interest). To enable proteomic analysis, we incorporated an affinity tag (e.g. ALFA tag) and also flanked each kinase substrate motif in the MKS with arginine residues, ensuring that trypsin digestion during sample preparation would result in kinase substrate peptides with distinct masses readily identifiable and quantifiable via MS^18^. The experimental workflow involves transfecting cells in culture with plasmids expressing the ProKAS polypeptide with kinase sensors and targeting elements of choice, after which cells are treated with a specific stimulus (Figure 1C). Following cell lysis and affinity purification, tryptic digestion generates a mixture of peptides and phosphopeptides that are quantified by MS to assess level of phosphorylation of each substrate sensor motif. The use of barcodes allows the generation of libraries of ProKAS biosensors and multiplexed analyses where each code can be linked to a specific targeting element, enabling analysis with spatial resolution (Figure 1D). Alternatively, barcodes can be linked to other variables, such as distinct drug treatments or genotypes, enabling comprehensive profiling of kinase signaling networks and facilitating high-throughput screens for drug effects or genetic perturbations.

**Figure 1.**
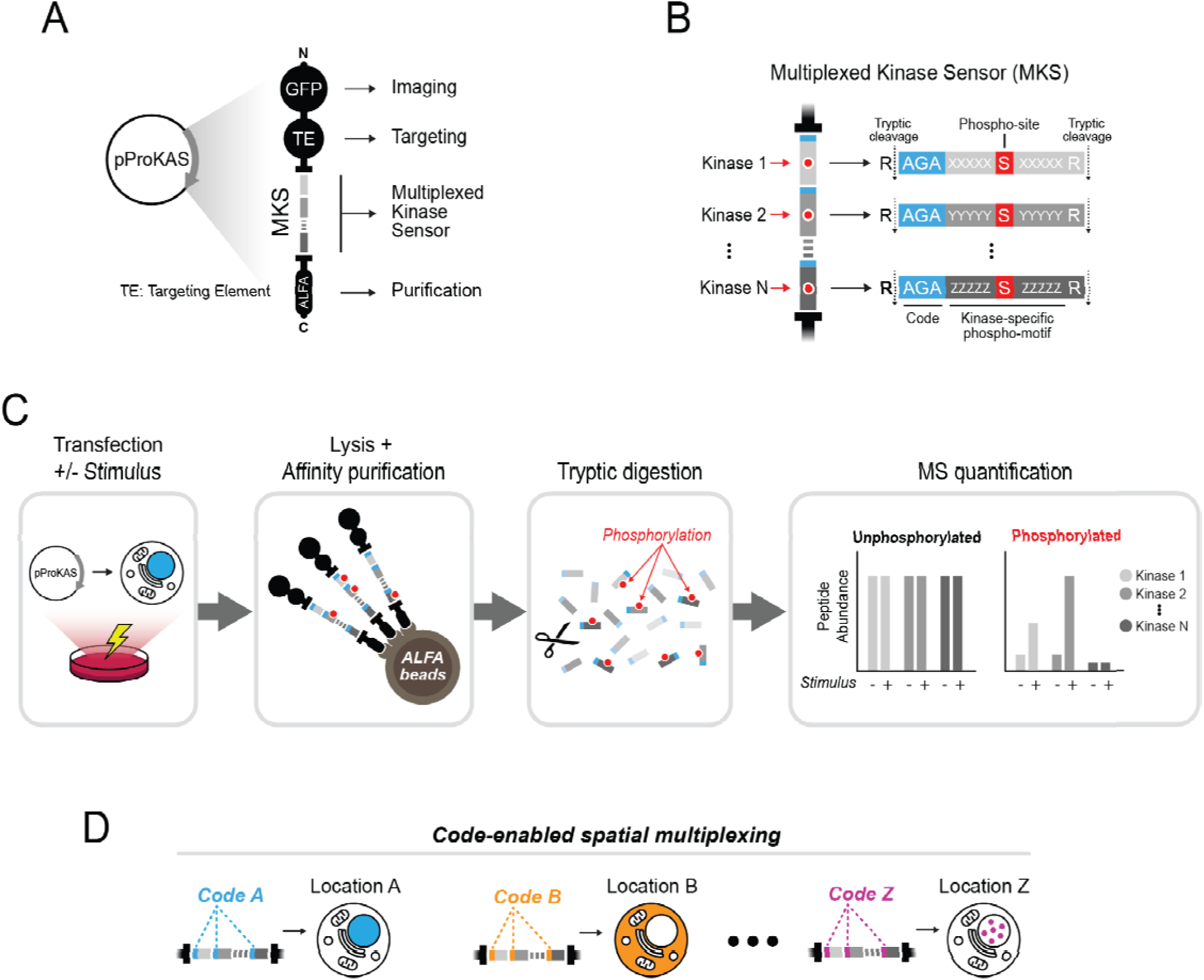
Design and rationale of ProKAS, a modular platform for multiplexed analysis of kinase activity using mass spectrometry. A) Schematic representation of the ProKAS construct for expression of the biosensor polypeptide containing fluorescent tags for visualization, a targeting element (TE) for subcellular localization, a multiplexed kinase sensor (MKS) module for detecting kinase activity, and an ALFA tag for high-affinity purification. B) Detailed view of the MKS module consisted of multiple kinase sensors, each featuring a kinase substrate motif, a barcode, and a flanking arginine residue allowing for tryptic digestion and independent MS-based quantification. C) Overview of the ProKAS workflow, starting with expression of biosensors via plasmid transfection. Expressing cell populations are then treated with a kinase stimulus of choice, after which ProKAS biosensors are purified from cell lysates, digested with trypsin, and the individual kinase sensors are quantified via MS in both their unphosphorylated and phosphorylated forms. D) Application of ProKAS for multiplexed spatial analysis of kinase activity. Multiple plasmids with distinct TEs matched to a specific barcode are co-transfected. MS analysis distinguishes the sensors based on the barcode mass allowing matching signal intensity of the specific peptide-probe to the respective cellular location.

### Generation of ProKAS sensors based on endogenous kinase substrates

The success of ProKAS relies on high-quality sensors used in the MKS module, wherein each peptide substrate must report the activity of a given kinase with sensitivity and specificity. We reasoned that the sequence of the substrate peptides could be based on ∼10-15 amino acid residues flanking phosphorylation sites on endogenous substrates, which could provide enough specificity for preferential targeting by a KOI. Phosphoproteomic data from large-scale experiments using kinase inhibitors or loss-of-function mutations would provide initial lists of phosphorylation events uniquely dependent on the KOI that are also compatible with MS detection. Once cloned into ProKAS, experiments should validate: (1) the ability of the MS to detect the phosphorylated peptide sensor; (2) the stimulus-dependent regulation of the probe phosphorylation; and (3) specificity of the probe phosphorylation by the KOI (Figure 2A).

**Figure 2.**
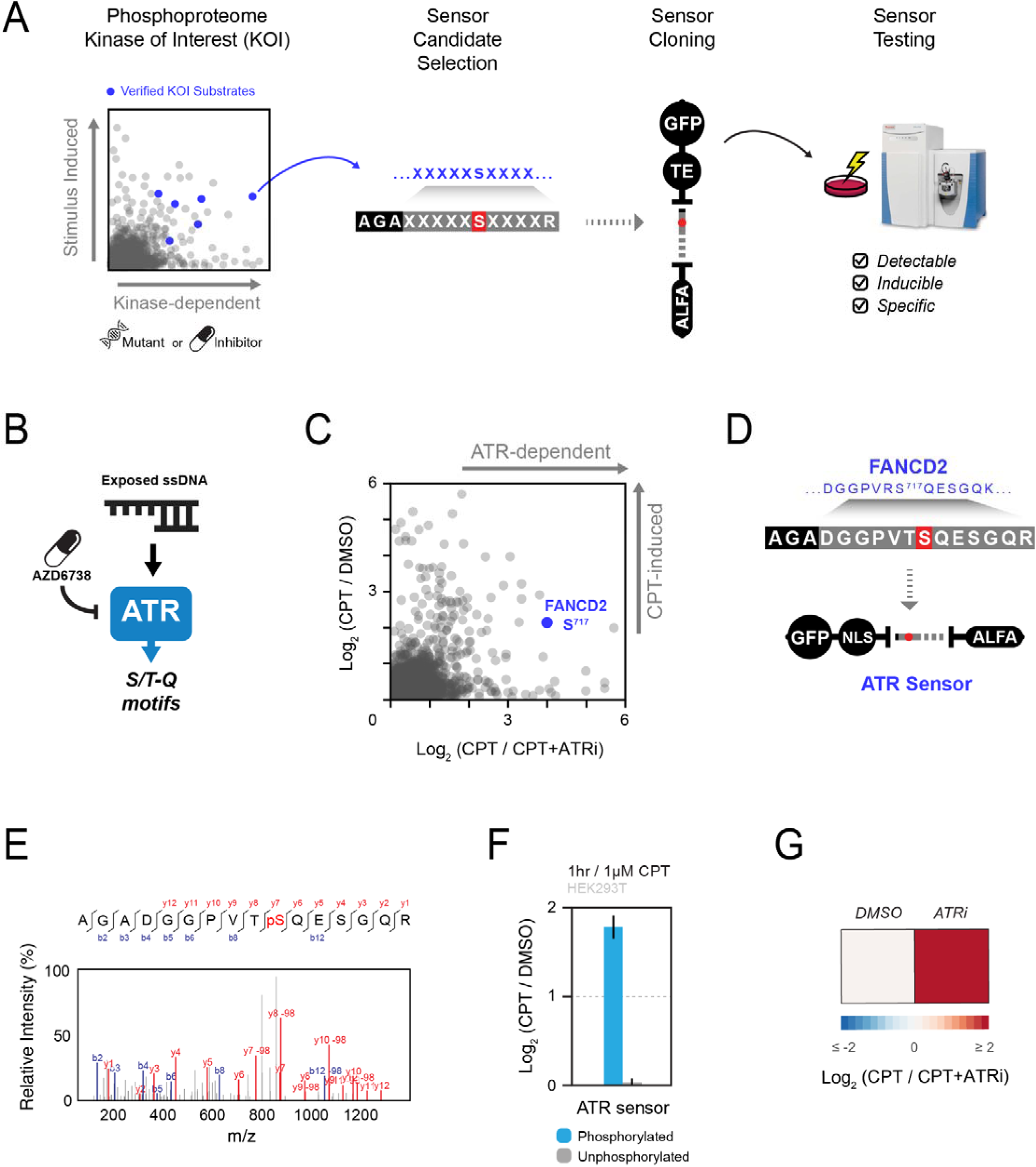
Development and validation of a ProKAS sensor specific for ATR using phosphoproteomic data. A) Schematic outlining of the strategy for designing kinase sensors based on phosphoproteomic data and known endogenous substrates. 10-15 sequences surrounding phosphorylation events detected to be dependent on a kinase of interest (KOI) are cloned into ProKAS constructs, which are then expressed in cells to test for kinase sensor detectability, inducibility, and specificity. B) Diagram showcasing that ATR is activated by single-stranded DNA damage, after which it preferentially phosphorylates substrates at the S/T-Q motif. Selective ATR inhibitors, such as AZD6738, are used in phosphoproteomic analyses to determine ATR-dependence of each detected phosphorylation event. C) Identification of FANCD2 S717 as an ATR-specific phosphorylation site that is also induced by genotoxin camptothecin (CPT) through phosphoproteomic analysis. D) Cloning of FANCD2 S717 as an ATR kinase sensor candidate into a ProKAS biosensor featuring a nuclear localization signal. E) MS/MS spectrum gathered for phosphorylated ATR sensor, showcasing successful detection of this sensor candidate by MS. F) MS analysis showing inducibility of phosphorylation of the ATR sensor upon treatment with 1 micromolar CPT. Unphosphorylated version of the sensor showed no significant change in abundance. G) MS analysis confirming specificity of ATR sensor phosphorylation using an ATR inhibitor. Error bars in F indicate the mean and standard deviation of triplicate independent experiments.

As a proof of principle, we generated a ProKAS construct to monitor the activity of the DNA damage signaling kinase ATR (Figure 2B). We followed the pipeline depicted in Figure 2A to identify specific ATR substrates suitable for a ProKAS sensor. Large-scale phosphoproteomic analyses of camptothecin (CPT)-treated cells, in the presence or absence of the ATR inhibitor AZD6738 (ATRi) were performed in pursuit of ATR-specific phosphorylation sites in the human proteome that could serve as templates for ATR sensors in the ProKAS biosensor. These experiments revealed dozens of phosphorylation sites in established ATR substrates that were strongly impaired by the inhibitor, indicating high specificity for ATR and no predominant targeting by related kinases ATM or DNA-PKcs. This analysis confirmed serine 717 of FANCD2, a previously known ATR substrate, as a phosphorylation site highly dependent on ATR that was also readily detectable via mass spectrometry ^19^ (Figure 2C). We therefore selected the segment encoding the 13 amino acids surrounding the FANCD2 S^717^ phosphosite and designed the ATR sensor by also adding a barcode and flanking R residues. This design results in the generation of a phosphopeptide slightly different from the endogenous FANCD2 after trypsin digestion. We then cloned this sequence into a ProKAS vector bearing a nuclear localization signal (NLS) as the targeting element, with the goal of monitoring ATR activity in the nucleus (Figure 2D). After transfection into HEK293T cells and treatment with CPT, the ATR ProKAS sensor was pulled down with anti-ALFA beads, trypsin digested, and analyzed by MS. PRM quantitation was used to directly compare phosphorylated sensors from SILAC-labeled cells treated (heavy R/K isotopes) and untreated (light R/K isotopes) with CPT, revealing that upon genotoxin treatment the abundance of phosphorylated ATR sensor is significantly increased, while its unphosphorylated form remains unchanged (Figure 2E). Confirming specificity, treatment with ATRi prior to CPT addition reduced the sensor phosphorylation to levels observed in the absence of genotoxin (Figure 2G). Overall, these results validate the strategy of extracting small (10-15 amino acids) sequences from endogenous substrates to design a ProKAS sensor that exhibits high specificity for a given KOI.

### *De novo* generation of a kinase-specific sensor peptide for ProKAS

We also developed a computational approach for the design of specific kinase probes by leveraging a dataset derived from positional scanning peptide array (PSPA) analyses of 303 human Ser/Thr kinases ^20^. The pipeline shown in Figure 3 scans PSPA-based kinase preference scores in a space of more than 500 billion 10-residue sequences to identify those exhibiting high specificity for a KOI, while minimizing cross-reactivity with the broader kinome (Figure 3A). Selected sequences are further evaluated based on a set of predicted features such as physicochemical properties, mass spectrometry retention time, and fragmentation patterns. After selection of 10 sequences with highest predicted specificity, lowest cross-reactivity and optimal properties, these are then cloned and expressed into one single ProKAS screening polypeptide for experimental validation (Figure 3B). This allows for the identification of the sequence motifs that are detectable in the MS and that demonstrate the best kinase inducibility and specificity, ultimately guiding the selection of the top KOI-specific peptide substrate sensor to be used for ProKAS applications.

**Figure 3.**
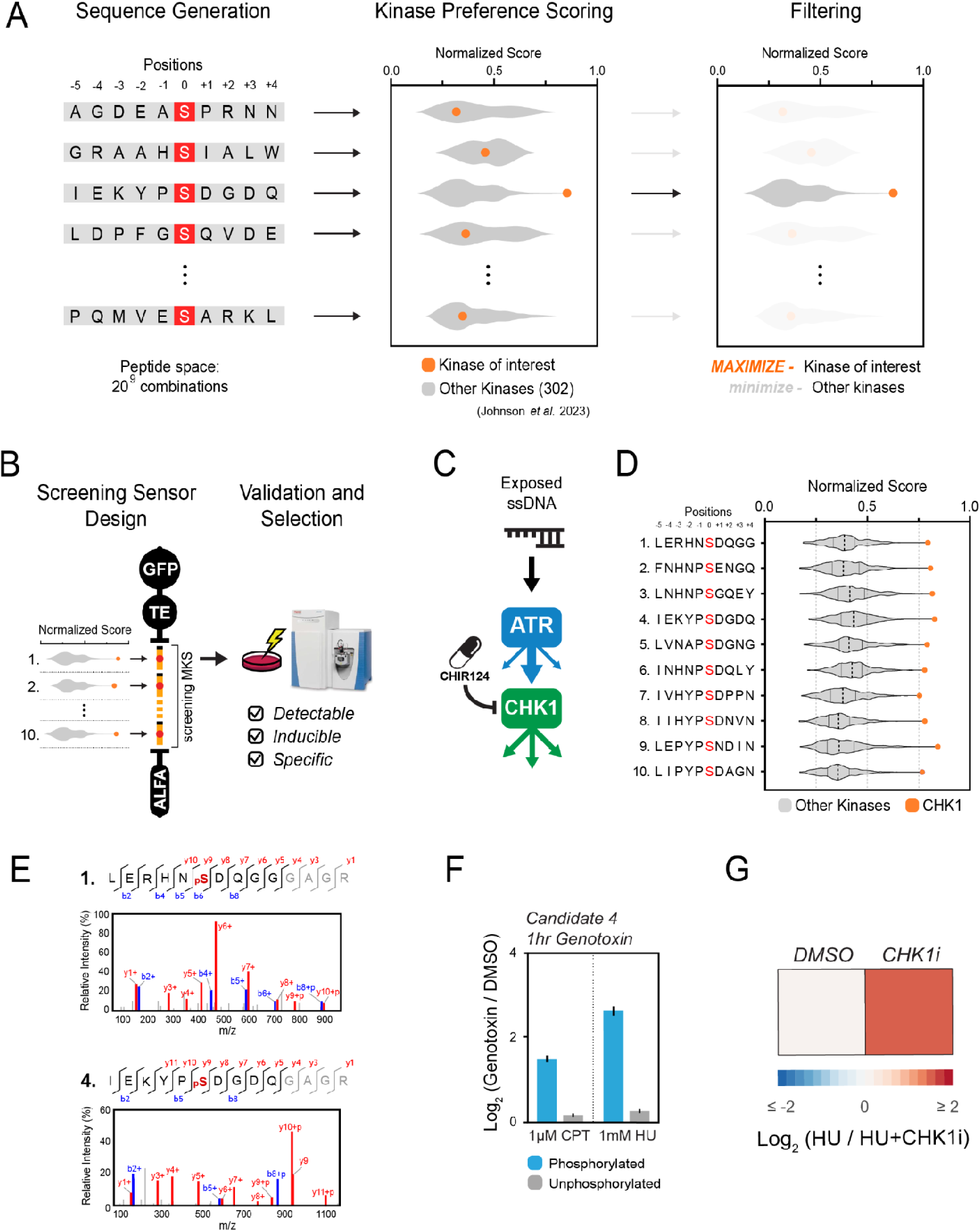
Pipeline for the computational design and experimental validation of a CHK1-specific ProKAS sensor. A) Overview of the computational approach for generating kinase-specific motifs based on PSPA data. B) Workflow for multiplexed screening and selection of MS-detectable kinase sensors from candidates generated and filtered *in silico*. C) Schematic illustrating CHK1 as a downstream effector kinase of ATR, and that CHK1 can be selectively inhibited by CHIR124. D) Selection of 10 candidate sequences for a CHK1 sensor based on PSPA scores as shown in (A). E) MS/MS spectra gathered for the phosphorylated form of the indicated CHK1 sensor candidates. Screening sensor with the candidates indicated in (D) were expressed in HEK293T cells treated with CPT. F) MS analysis showing inducibility of phosphorylation of the CHK1-4 sensor candidate upon genotoxin treatment. G) MS analysis confirming specificity of CHK1-4 sensor phosphorylation using a CHK1 inhibitor. Error bars in F indicate the mean and standard deviation of triplicate independent experiments.

We applied this pipeline to develop a probe for CHK1, a key downstream effector kinase activated by ATR in response to DNA damage^21^ (Figure 3C). While CHK1 is known to preferentially phosphorylate substrates containing Arg or Lys residues at the position -3 relative to the phosphosite, the PSPA data confirmed this preference and further revealed that CHK1 exhibits strong preferences for bulky hydrophobic residues at the position -5 (e.g., Leu, Iso, and Phe), as well as a moderate preference for Pro and Asn at the position -1 ^20,22–24^. We selected sequences exhibiting high PSPA-bases scores for CHK1, while minimizing scores for the other 301 kinases (Figure 3D). These contained Arg/Lys at position -3 but also incorporated variations at this and other positions to explore the impact on CHK1 specificity and sensor performance. A CHK1 ProKAS screening sensor was generated with a MKS module containing 10 candidate peptide sensors and a NLS targeting element to direct the sensor to the nucleus. MS analysis could readily detect tryptic peptides for all 10 candidate sensors (SD5). However, upon genotoxic stress, only two phosphorylated peptide substrates were identified, CHK1-1 (LERHNSDQGGGAGR) and CHK1-4 (IEKYPSDGDQGAGR), both containing Arg or Lys at the position -3 (Figure 3E). The CHK1-4 peptide, selected as our final CHK1 sensor, demonstrated high inducibility upon cell treatment with both CPT and HU (Figure 3F). The greater induction observed under HU treatment is consistent with the known effect of this drug in generating long stretches of ssDNA, which robustly activates ATR and consequently CHK1, whereas CPT causes DSBs and thereby predominantly activates ATM ^13,25^. We confirmed that the CHK1-4 sensor was specifically phosphorylated by CHK1 through the use of a selective CHK1 inhibitor CHIR-124 ^26^(Figure 3G). Overall, these results confirm that Arg/Lys at the position -3 is a strong determinant of CHK1 specificity, and are consistent with the *in vitro* specificity screen assay showing that Leu/Iso at the position -5 favors CHK1 phosphorylation recognition *in vivo*. Importantly, in addition to generating preferential sequences for phosphorylation by the kinase of interest, this strategy is expected to also generate sequences that are less prone to be targeted by other kinases. The fact that phosphorylation of the sensor was impaired by CHK1 inhibitor supports the notion that the sensor is not being predominantly targeted by another kinase, even constitutively. These findings also highlight the need for an experimental screening step given the majority of *in silico* generated sequences did not behave as effective sensors for our kinase of interest.

### Multiplexed monitoring of ATR, ATM, and CHK1 activity with ProKAS

We next assessed the use of the ProKAS platform for monitoring the activity of multiple kinases simultaneously. We generated a multiplexed “DDR” ProKAS sensor to simultaneously monitor the activity of ATR, CHK1 and ATM kinases (Figure 4A-B). For the ATM ProKAS sensor, we employed the phosphoproteome-guided strategy, revealing a phosphorylation event in the known ATM substrate 53BP1, which was ultimately selected as our sensor peptide (Figure S1). The DDR ProKAS containing a tandem array of the ATR, ATM and CHK1 sensors (Figure 4B) was expressed in cells that were mock treated or treated with either CPT or HU to predominantly generate DSBs or stalled forks, respectively ^27,28^. We first confirmed that swapping substrate peptide positions within the MKS module did not alter their level of inducibility upon genotoxin-induced kinase activation (Figure S2). We next assessed if each of the sensors retained the specificity for the respective KOI. As shown in Figure 4C, the use of specific kinase inhibitors confirmed that the sensors within the context of the ProKAS peptide array were specifically phosphorylated by the expected kinase. Inhibition of ATR under both genotoxin conditions revealed a reduction in the phosphorylation of the CHK1 sensor, consistent with the activation of CHK1 being canonically controlled by ATR ^13,29^. Inhibition of CHK1 alone did not reduce the phosphorylation of ATR or ATM kinase sensors in cells treated with CPT. However, phosphorylation of the ATM sensor increased in HU-treated cells upon CHK1 inhibition, consistent with CHK1 playing key roles in preventing the collapse of stalled replication forks in HU, which causes DSBs and ATM activation ^30^. As expected, ATM inhibition robustly and specifically ablated phosphorylation of the ATM sensor upon CPT treatment, and also displayed some level of inhibition of the ATR and CHK1 probes due to likely effects in inhibiting DNA end resection during the DSB response ^31,32^. We also confirmed that expression of these kinase sensors did not impair the endogenous DNA damage response by blotting for markers of ATR and ATM activation after treatment with CPT and HU (Fig S3)^33,34^.

**Figure 4.**
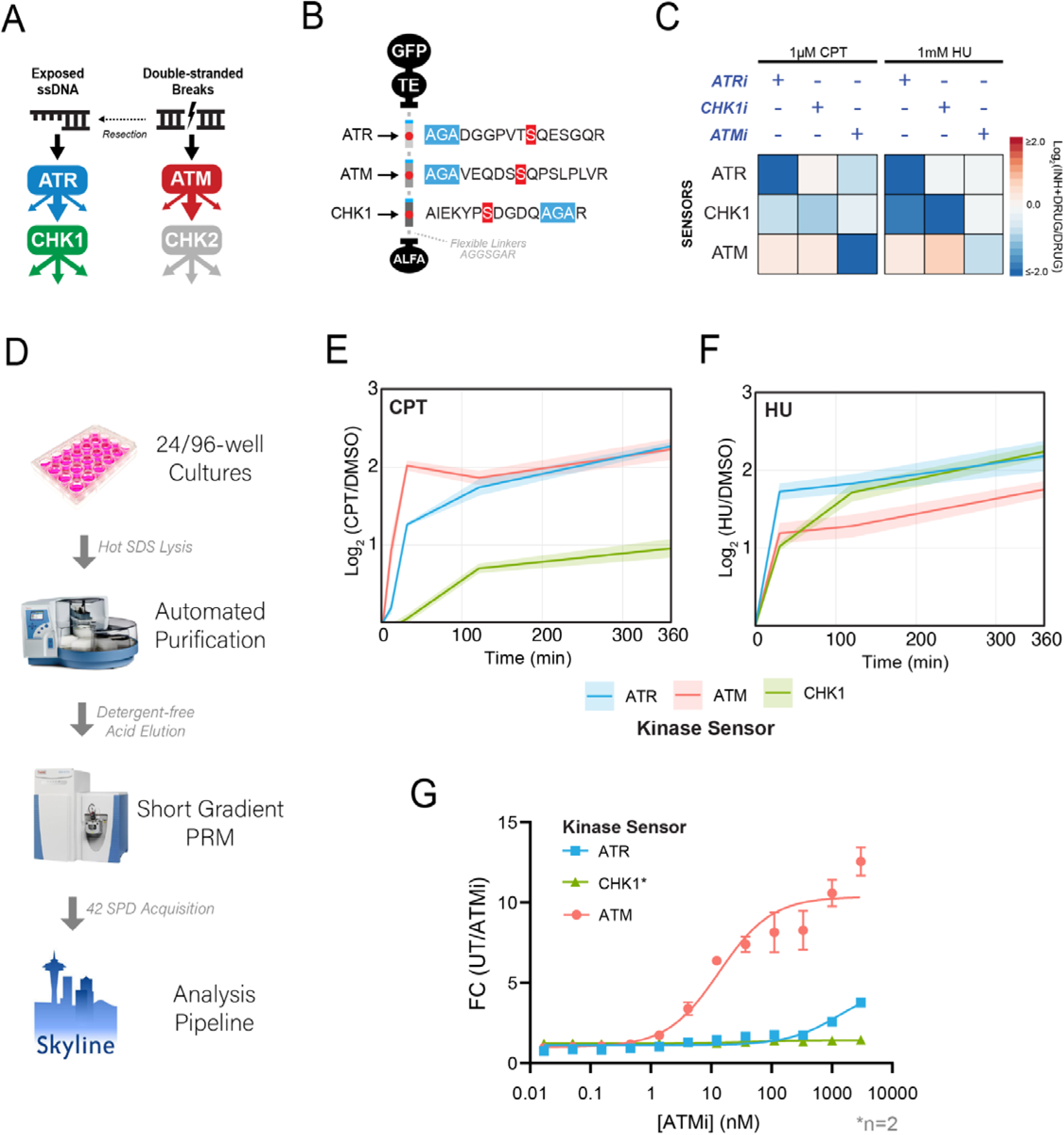
Multiplexed analysis of DDR kinase activities using ProKAS. A) Schematic illustrating the canonical mechanisms of activation for DDR kinases ATR, ATM, and CHK1. B) Design of a triplexed ProKAS construct containing kinase sensors for ATR, ATM, and CHK1. C) Validation of kinase specificity of each sensor within the triplexed ProKAS construct via treatment with selective kinase inhibitors for each KOI in the presence of either 1 micromolar CPT or 1 millimolar HU. ATR, ATM, and CHK1 were inhibited with 1 micromolar AZD-6738, 50 nanomolar AZD-0156, and 500 nanomolar CHIR-124, respectively. D) Flowchart showing primary components of the semi-automated high throughput pipeline enabling larger scale ProKAS experiments. E) MS analysis showing kinase activation dynamics in response to 1 micromolar CPT over 6 hours in HEK293T cells expressing the triplexed ProKAS construct. F) MS analysis showing kinase activation dynamics in response to 1 millimolar HU over 6 hours in HEK293T cells expressing the triplexed ProKAS construct. G) MS analysis showing inhibitor titration using ProKAS to demonstrate the potency and selectivity of ATM inhibitor AZD0156. Cells were treated with 1 micromolar CPT for 30 minutes to activate ATM. Error bars/envelopes in E, F, and G indicate the mean and standard deviation of triplicate independent experiments, except for CHK1 sensor quantification in G which comprises duplicate independent experiments.

In addition to multiplexing, other advantages of ProKAS include the throughput and highly quantitative nature of the analyses, which should allow for precise kinetic studies on kinase signaling. We implemented a semi-automated pipeline for processing up to dozens of samples in 24-well cell culture plates (Figure 4D), which enabled temporal kinetic analysis of ATR, ATM, and CHK1 signaling in response to CPT and HU (Figures 4E-F). Consistent with the expected behavior of these kinases upon treatment with CPT, which induces DSBs, ATM showed rapid and robust activation within 10 minutes, reflecting its role as a primary responder to DSBs. ATR activation followed, ultimately matching ATM activation levels at later timepoints, consistent with a need for longer times for DNA ends to be resected and to support ssDNA-mediated ATR activation^35^. The slower kinetics of CHK1 activation was consistent with it being a kinase downstream of ATR. In contrast, HU treatment revealed distinct kinetics of activation for these DDR kinases. Consistent with HU causing the stalling of replication forks and rapid ssDNA exposure, ATR was rapidly activated, whereas the ATM activation never reached levels as high as ATR activation (Figure 4F). CHK1 phosphorylation was slower initially, but ultimately surpassed ATM. These observations align with the canonical behaviors of these kinases in response to DNA damage and highlight the ability of ProKAS to generate highly quantitative data on kinetics of kinase activation. Finally, we leveraged the multiplexed and throughput capabilities of ProKAS to assess the inhibitory potency of an ATM-specific inhibitor (AZD0156) *in vivo*. We performed an inhibitor titration experiment in HEK293T cells expressing the triplexed DDR ProKAS sensor, monitoring the phosphorylation levels of all three sensors simultaneously. As expected, AZD0156 potently inhibited ATM activity, with an IC50 of 9 nanomolar, closely mirroring the reported value obtained using *in vitro* assays with purified enzymes ^36^ (Figure 4G). Importantly, the inhibitor exhibited high selectivity, showing no significant effect on CHK1 activity and minimal impact on ATR activity even at high concentrations (IC50 > 5 micromolar). This experiment showcases the power of ProKAS in enabling precise, multiplexed, and *in vivo* assessment of kinase inhibitor efficacy and selectivity, offering a valuable tool for drug discovery and development.

### Spatial analysis of kinase signaling using location-barcoded ProKAS

In addition to its quantitative and multiplexed capabilities, ProKAS was designed to also enable the spatial analysis of kinase activity in cells. As shown in Figure 1D, the ProKAS construct features a targeting element that controls cellular localization and that can be linked to a specific tri-amino acid barcode embedded within each sensor peptide. Upon co-transfection of cells with ProKAS constructs containing different targeting elements, detection of distinct barcoded peptides by MS should reveal the relative levels of kinase activity in different cellular locations. As a proof of concept, we used a nuclear localization signal (NLS) or a nuclear export signal (NES) as a targeting element to direct the ProKAS polypeptide to the nucleus or cytosol, respectively (Figures 5A-B). The AGA barcode was used in the construct containing the NLS and a GAG barcode was used in the construct with the NES. Importantly, addition of different barcodes (AGA, GAG or GVG) to the substrate sensor peptides within the MKS module did not alter their efficiency of phosphorylation upon kinase activation (Figure S4).

**Figure 5.**
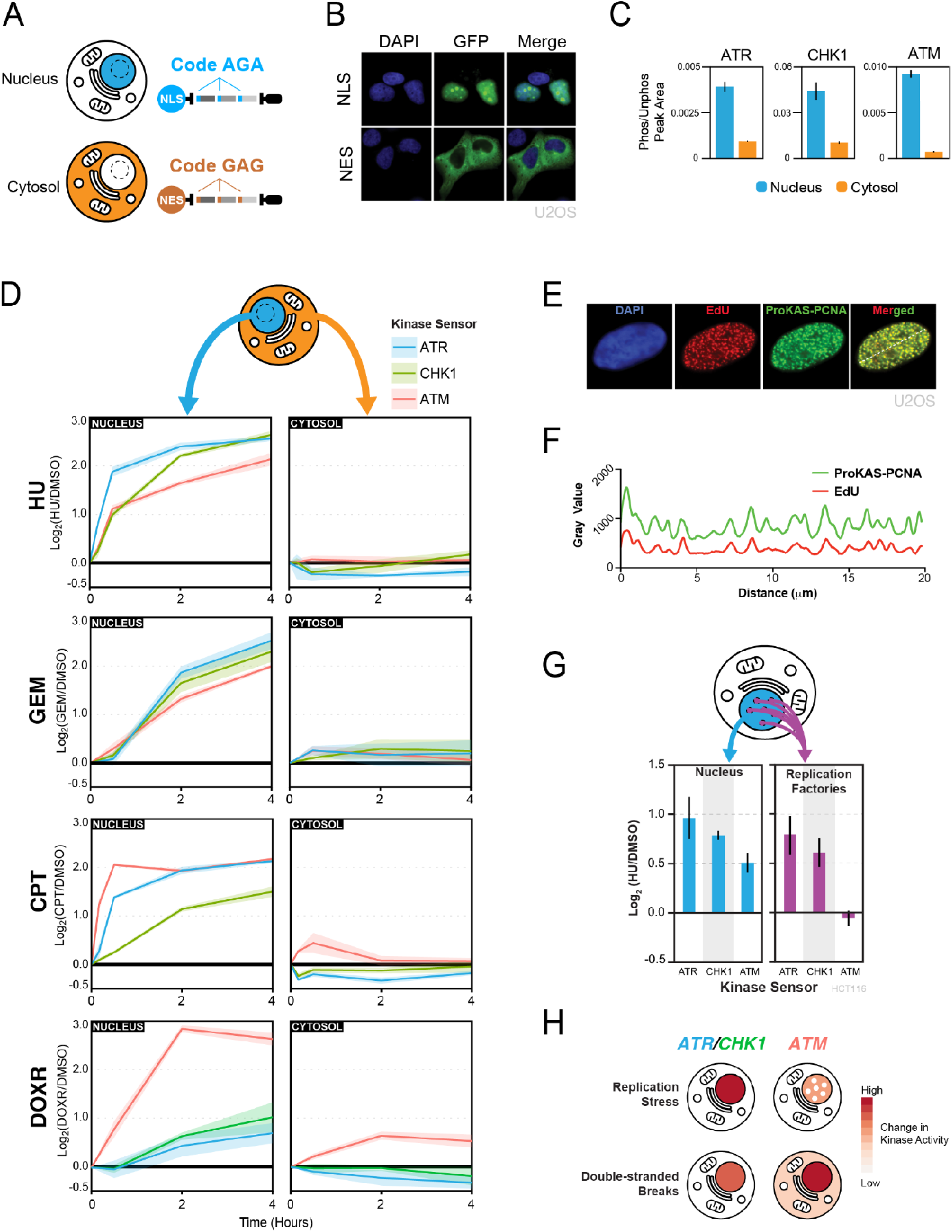
Spatially-encoded ProKAS for analysis of DDR kinase activities. A) Illustration of nuclear and cytosolic ProKAS biosensors featuring distinct codes for simultaneous monitoring of DDR kinase activities in both locations. B) Microscopy illustrating the localization of ProKAS biosensors containing an NLS or an NES. C) Peak area ratios for DDR kinase sensors showing the proportion of phosphorylated sensor peak area to unphosphorylated peak area. Data based on MS analysis of cells not treated with any genotoxin. D) MS analysis of ProKAS sensors simultaneously monitoring DDR kinase signaling kinetics in both the nucleus and cytosol in response to various genotoxic agents. GEM and DOXR were performed label free. E) Microscopy showing the co-localization of the ProKAS biosensor (utilizing PCNA as a targeting element) with EdU foci. F) Densitometry analysis of EdU and ProKAS-PCNA signal across the white line drawn across the nucleus in panel E, showing signal coincidence between the EdU and ProKAS-PCNA foci formed. G) MS analysis simultaneously monitoring the effect of HU on ProKAS biosensors containing an NLS or PCNA as the targeting element. H) Schematic illustration of different spatial distributions of ATR, CHK1, and ATM kinase activity observed upon replication stress or DSBs. Error bars/envelopes in C, D, and F indicate the mean and standard deviation of triplicate independent experiments.

ATR, ATM, and CHK1 are DNA damage signaling kinases known to operate primarily inside the nucleus. While cytosolic roles for these kinases have been proposed ^37–39^, potential non-nuclear functions remain elusive due, in part, to the need of more quantitative tools capable of monitoring the activity of these kinases with rigorous specificity and spatial resolution. Using ProKAS, we analyzed cells co-expressing nuclear and cytosolic versions of the multiplexed ATR, ATM and CHK1 sensors differentially barcoded based on their location (Figures 5C-E). The intensity of the phosphorylated peptide in each compartment was divided by the intensity of the corresponding unphosphorylated peptide in that compartment, yielding normalized values of peptide phosphorylation, a rough measure of stoichiometry of phosphorylation. As shown in Figure 5C, while we were able to detect phosphorylation of all three sensors in both nucleus and cytosol in the absence of any genotoxic treatment, the constitutive levels of kinase sensor phosphorylation were significantly lower for the cytosol-localized peptide sensors compared to their nucleus-localized counterparts. Such high levels of constitutive kinase signaling in the nucleus are consistent with the high endogenous levels of DNA replication stress in HEK293T cells^19^. We next monitored the kinetics of ATR, ATM and CHK1 signaling in the nucleus and in the cytosol following treatment with replication stress and DNA damage inducing agents (Figure 5D). Each of the four genotoxic agents exhibited distinct patterns of kinase activation over time, highlighting the distinct modes through which these agents perturb DNA replication and/or cause DNA breaks. For example, while HU induced a rapid increase in nuclear ATR signaling, gemcitabine (GEM) induced a delayed response in which the nuclear activity of all three kinases slowly increased with similar kinetics. In contrast, CPT was marked by an ultra-fast increase in nuclear ATM signaling followed by a very fast increase in nuclear ATR signaling that reached similar levels as the ATM signaling. Doxorubicin (DOXR) was drastically different, inducing a strong increase in nuclear ATM signaling that was largely uncoupled from ATR-CHK1 signaling. Interestingly, while HU and GEM did not result in any significant changes in ATR, ATM or CHK1 kinase activity in the cytosol, ATM activity was specifically detected in the cytosol when treating cells with CPT or DOXR, both of which are agents that induce DSBs (Figure S5). ATM has been shown to localize to various cytosolic locations in specific situations ^40,41^, but its level of activation outside the nucleus has been difficult to measure. While the increase in cytosolic ATM signaling was relatively mild compared to the detected increase in nuclear ATM signaling, it is important to note that the larger volume of the cytosol is likely diluting the observed induction, especially if the source of the non-nuclear ATM activation is at a specific cellular location, such as the external membrane of a specific organelle. Since we did not detect an increase in cytosolic ATR or CHK1 signaling in response to any of the treatments tested, the increase in cytosolic ATM signaling induced by DSB is likely a specific feature of ATM activation or signaling propagation, and is unlikely to represent a non-selective leak of nuclear kinases into the cytosol.

To demonstrate the ability of ProKAS to probe differences in kinase activity within nuclear sub-compartments, we focused on sites of DNA synthesis. Upon HU treatment, sites of DNA synthesis are expected to accumulate stalled replication forks and single-stranded DNA, resulting in the selective induction of ATR-CHK1 signaling^15^. We localized the ProKAS biosensor to sites of DNA synthesis by using the processivity factor PCNA clamp, a key component of replication forks, as the targeting element. Microscopy analysis confirmed that the PCNA-containing biosensors co-localized with nascent DNA, as visualized via incorporation of EdU (Figure 5E, F). Using HCT116 cells, we then co-expressed ProKAS constructs having PCNA-NLS or only NLS as the targeting element, allowing comparison of kinase signaling diffused in the nucleoplasm with signaling localized to sites of DNA synthesis. Upon treatment with 1 millimolar HU for 1 hour, biosensors localized to the nucleus with an NLS alone showed induced phosphorylation of all three kinase sensors (Figure 5G). In contrast, the ATM sensor in the PCNA-containing ProKAS construct exhibited no induced phosphorylation upon HU treatment, while the ATR and CHK1 sensors were robustly induced. This finding is consistent with the notion that fork stalling specifically activates ATR-CHK1, but not ATM, which mainly responds to DSBs ^13,14^. This finding also shows that while HU does result in ATM activation, the nature and location of such activation is different than where ATR-CHK1 are activated, and is unlikely to occur at stalled forks containing PCNA. Overall, these results demonstrate the ability of ProKAS to monitor kinase signaling with spatial resolution and define key differences in the cellular location of ATR and ATM signaling upon replication stress and DSBs (Figure 5H).

## Discussion

In the last decades, technological advances such as genetically encoded fluorescent biosensors, analog-sensitive engineered kinases, and MS-based phosphoproteomics have revolutionized our ability to study kinase action in cells ^1,6,9,39,42^. Yet, much remains unknown about how kinases act in cells to regulate and coordinate cellular processes and responses. A major obstacle is the lack of versatile and generalizable techniques capable of systematically integrating quantitative, spatial, and temporal analyses of kinase action.

The ProKAS system introduced here expands the set of tools available for monitoring kinase activity in living cells. ProKAS overcomes some of the limitations associated with traditional fluorescence-based kinase sensors by integrating the strengths of MS with the versatility of peptide-based biosensors. Many fluorescence-based kinase sensors involve elaborate engineering that is constrained not only by the identification of substrate peptide sequences that are specific to the KOI, but also by the requirement that, upon phosphorylation, the sequences are recognized by a phosphorylation-binding domain (e.x. FHA domain for FRET sensors) or induce changes in subcellular localization (e.x. nucleus to cytosolic translocation) ^7^. A major advantage of ProKAS is the flexibility in the generation of the sensor sequences, which are constrained mainly by the KOI specificity and by the ability to be detected by MS. Since sensors can be designed based on phospho-peptides detected in phosphoproteomic datasets, the requirement of MS-detectability is already satisfied upon phosphopeptide selection. In the case of the *in silico* designed sensors, our experience is that most sequences of around 10 amino acids can be detected by MS, allowing flexibility in sequence selection, which becomes mainly dictated by kinase specificity. We expect that the reduced stringencies in sequence design compared to FRET sensor design should make ProKAS a more generalizable tool for exploring the action of the broader kinome. We also observed higher dynamic ranges in the recorded ProKAS sensors compared to FRET sensors. For example, the recently reported FRET sensor for CHK1 showed a lower than 2-fold change in signal upon replication stress (typical of FRET sensors), whereas our CHK1 sensor displayed over 4-fold increase in phosphorylation upon replication stress ^43^.

Another important feature of ProKAS is the ability to monitor multiple kinases simultaneously (through the use of tandem sensor arrays), which is expected to be essential for studying complex signaling responses and for high-throughput analyses. In addition to having multiple sensors in one ProKAS construct, the use of amino acid barcodes further expands the possibilities of studying kinase signaling, especially for spatially resolved analyses. We demonstrated these multiplexing features of ProKAS by generating a multi-sensor module for the DNA damage response kinases ATR, ATM, and CHK1 and by capturing their activity over distinct cellular locations. We showed that through the use of specific targeting elements, ProKAS can resolve sub-organellar locations, such as sites of DNA synthesis within the nucleus. In these spatially resolved analyses, since the locations are pre-determined by validated targeting elements, data analysis and interpretation are straightforward. In the future, we expect that generation of libraries of targeting elements for a range of specific cellular locations will allow the systematic spatial analysis of the action of multiple kinases simultaneously. Barcoding can also be further expanded through the use of 4 amino acid barcodes to enable the multiplexed analysis of dozens of locations. The tandem sensor arrays may also accommodate additional 10-15 amino acid sensor sequences to monitor, in principle, dozens of kinases at once. The major factor limiting the number of sensors is likely the size of the ProKAS construct, as very large constructs would likely lead to reduced protein expression level. Removal of GFP or optimization of linker sequences in the current version of ProKAS would free up additional space to accommodate dozens of sensors. We also demonstrated that the ProKAS system can be employed in high throughput pipelines through semi-automated cell lysis, affinity capture, and sample preparation, allowing analyses of samples from 24-well plates on a QE-HF MS instrument. From cell treatment to quantitative readout, the experimental pipeline could often be completed in less than two days. We predict that the use of more sensitive MS systems should allow the use of 96-well plates, ultimately enabling platforms for screening the impact of drugs on kinase signaling.

Of importance, our work demonstrated the feasibility of generating sensors that distinguish the specificities of closely related kinases, such as ATR and ATM, which are known to have a similar preference for S/T-Q motifs^44^. Interestingly, the specificities conferred by the 10 amino acid sequence surrounding the phosphorylation site of the ATR or ATM sensors we generated here were based on phosphoproteomic data and are not predicted to be highly specific according to existing *in vitro* preference data ^20^. Since there may be yet unknown levels of combinatorial effects within these amino acid sequences, we expect that sequence information embedded within the 10-15 amino acids surrounding the phospho-acceptor site on the substrate might be sufficient to determine specificity toward a unique kinase, and that such sequence determinants can be exploited in kinase sensor design. In this direction, the sensor design process may be improved by incorporating machine learning approaches trained on large datasets of experimentally validated substrates. In any case, we expect future work to expand ProKAS to other kinases.

The application of ProKAS to the study of ATR, ATM and CHK1 provided new insights into how these kinases respond to different genotoxic drugs. The quantitative kinetic data defined major differences in how different modes of replication stress and DNA damage promote differential profiles of activation for each of these kinases. We foresee that systematically expanding these analyses to large panels of drugs will aid in understanding the mechanisms of drug action with greater detail. It will also be important to expand the analyses to panels of cancer cells as well as untransformed cells to define how the impact of genotoxic drugs changes depending on cell types and oncogenic state. We also detected a cytosolic mode of ATM signaling induced by drugs that cause DSBs. Further work will be necessary to define the origin of this cytosolic signaling, if it originates in the nucleus and propagates to the cytosol, or if it originates at other organelles such as the mitochondria. Another possibility is that DNA fragments may leak from the nucleus or the mitochondria and activate a specific pool of cytosolic ATM molecules. Interestingly, ATM is known to have a non-canonical mode of activation triggered by oxidative stress, raising the possibility that the cytosolic ATM signaling we observed could be caused by some form of oxidative stress caused by CPT and DOXR ^45^. We predict that ProKAS will be useful to study the ATM response to oxidative stress with quantitative kinetics and spatial resolution.

In conclusion, ProKAS is a new tool for quantitative and multiplexed analyses of kinase action with spatial resolution. The possibilities of integrating ProKAS with high-throughput analyses positions this technology as a promising tool for systems biology research and drug discovery. Future developments in computational design, expanded substrate libraries, and integration with other omics technologies will further enhance the utility and impact of ProKAS in unraveling the complexities of kinase biology.

### Limitations

While ProKAS provides a flexible approach to monitoring kinase activity in living cells, it currently involves the overexpression of biosensors from plasmid libraries which in turn requires that cells effectively uptake and express exogenous DNA. While HEK293T and HCT116 cells are amenable to common transfection protocols, some cell lines may pose challenges for efficient transfection. Additionally, the biosensor is currently expressed at high levels in order to ensure high sensitivity of the technique. This has proven effective for locations of interest thus far, but for some subcellular locations like telomeres or centromeres, dramatic overexpression may lead to spatial oversaturation. To address this, attenuation of expression via promoter modifications can be performed, but at the expense of assay sensitivity ^46^. Additionally, targeting elements must be chosen with care so as to avoid cytotoxicity. Overcrowding specific subcellular loci may lead to changes in cell behavior, thereby causing aberrant signaling to be detected. Some proteins that may be desirable as targeting elements may also prove toxic to some cell types regardless of expression level. ProKAS also currently monitors the kinase activity in a cell population and, in contrast to FRET-based kinase probes, does not have the ability to generate data from single cells. Moreover, while the design of sensors was successful for the kinases featured in this study, the possibility remains that several kinases may require more than just the 10 flanking amino acid residues to recognize and phosphorylate a sensor sequence with specificity; for any such kinase, the ProKAS approach to sensor design would require consideration of additional variables like substrate structure, binding partners, docking sites, or prior post-translational modification of substrate sequences.

## Supporting information

de novo algorithm script

KingFisher program

Skyline document with settings for ProKAS experiments

Supplemental tables 1 through 5

Plasmid sequencing results

## Data Availability

Mass spectrometry data generated for this study are available through PRIDE, wherein phosphoproteomics data can be found under PXD identifier PXD059064 and targeted data under PXD059058 (https://www.ebi.ac.uk/pride/)^47^. Targeting element and MKS sequences are included in supplemental data (SD1 and SD2). Phosphoproteome results tables and a summary of CHK1 sensor candidate testing are also included in supplemental data (SD3, SD4, and SD5). Whole plasmid sequencing results for all ProKAS plasmids are included in .fasta format as part of the supplement. A Skyline document configured with all peptide and transition settings used is included as a supplemental file. A method file for execution of automated ALFA affinity purification on a KingFisher is also included.

## Code Availability

Python code and accompanying files for *in silico* generation of kinase sensor candidates are included in the supplement.

## Acknowledgements

We thank Beatriz S. Almeida for technical support and members of the Smolka Lab for valuable discussions. This work was supported by grants from the National Institute of Health, R35GM141159 and R01HD095296 to MBS, and F31CA281247 to WJC.

## Author contributions

Conceptualization, WJC, MVASN, MBS; Data Curation, WJC, MVASN, DVM, YR, MW; Formal Analysis, WJC, MVASN, DVM, YR, MW; Funding acquisition, WJC, MBS; Investigation, WJC, MVASN, DVM, YR, MW; Methodology, WJC, MVASN, MBS; Project Administration, MBS; Resources, MBS; Software, MVASN; Supervision, MBS; Validation, WJC, MVASN, DVM, YR, MW; Visualization, WJC, MVASN, MW, YW; Writing - original draft, WJC, MVASN, MBS; Writing - review and editing, WJC, MBS.

## Methods

### Plasmid Construction

All pMKS plasmids constructed in this study were generated using Gibson assembly using HiFi DNA Assembly Master Mix (NEB). PCR reactions were performed with primers ordered from Integrated DNA Technologies (IDT) and Q5 High-Fidelity 2X Master Mix (NEB). All gBlock synthetic gene fragments (targeting elements and MKS modules) were ordered from IDT and cloned into a plasmid containing EGFP under the control of a CMV promoter (Addgene plasmid 46957). Full plasmid sequences in .fasta format are available in Supplementary Data.

### Cell Culture and Transfection

HEK293T, U2OS, and HCT116 cells (ATCC) were cultured in DMEM (Gibco) supplemented with MEM Non-essential amino acids (Corning), Penicillin-Streptomycin (Sigma), and dialyzed Fetal Bovine Serum (Sigma). For SILAC cultures, DMEM for SILAC was used instead (Thermo Scientific), supplemented with either normal isotopes (“Light” channel) or heavy isotopes of Lysine and Arginine (“Heavy” channel). Isotopes of Lysine and Arginine used were L-Lysine-13C6,15N2 HCl and L-Arginine-13C6,15N4 HCl (Sigma). Cultured cells were passaged using 0.05% Trypsin-EDTA solution (Gibco). Reverse transfection of HEK293T cells with ProKAS plasmids was performed using 1 milligram per milliliter polyethylenimene and DMEM for SILAC to which no amino acids or serum was added (“SFS”). For a 12-well plate, 7.2ug of plasmid DNA was mixed with 900 microliters of SFS, after which 36 microliters of 1 milligram per milliliter PEI was added (5:1 ratio of PEI:DNA) (Thermo Scientific). This mixture was allowed to incubate at room temperature for 20 minutes. Cells were detached with trypsin and counted using a Cytosmart cell counter (Corning) before being suspended in SILAC media to which the mixture of PEI, plasmid, and SFS were added prior to plating. Cells were allowed to grow for 36-48 hours before treatment. HCT116 cells were transfected with a similar procedure substituting PEI with Mirus TransIT reagent (Mirus Bio). When using 6-, 24-, and 96-well plates, proportions of reagents were scaled with the surface area of plates and wells used.

### Manual Affinity Purification

Upon completion of cell treatments, media was aspirated from the plates and 150 microliters of modified radioimmunoprecipitation assay (mRIPA) lysis buffer containing 0.2% SDS was added to the cells before plates were placed on a hot plate set to 200□ for 10 seconds. The lysis buffer and cells were agitated before adding 450 microliters of mRIPA lysis buffer to dilute the SDS concentration below 0.1% SDS. Modified RIPA buffer was 50 millimolar Tris-HCl pH 8.0, 150 millimolar NaCl, 1% tergitol, and 5 millimolar EDTA. For SILAC experiments, lysates for both the light and heavy SILAC channels were then pooled in 1.5 milliliter microcentrifuge tubes, whereas for label-free experiments no pooling was performed. Lysates were then sonified using a Branson 1/8” probe tip sonifier for 5 seconds at 12% amplitude to shear genomic DNA and reduce sample viscosity. Sonified lysates were centrifuged for 5 minutes at 13,000G+ in a 4□ centrifuge. For each pooled lysate, 4 microliters of magnetic ALFA Selector ST affinity purification beads were equilibrated in Protein Lo-bind Eppendorf tubes containing 150 microliters of lysis buffer, after which the beads were immobilized via magnetic rack and the lysis buffer was aspirated. Centrifuged lysates were transferred to these tubes containing equilibrated beads, after which they were nutated at 4□ for 10 minutes. After incubation with the beads, the samples were placed in magnetic racks to immobilize the beads and the supernatants were aspirated. Beads were washed twice with 500 microliters of lysis buffer and once with 1 milliliter of sterile deionized water, with each wash involving 2 minutes of nutation at room temperature. ProKAS modules were eluted from the beads using 80 microliters of acidic elution buffer (0.1 molar Glycine HCl, 150 millimolar NaCl, pH 2.2) for 2 minutes at room temperature. Elutions were transferred to new Lo-Bind tubes containing 20 microliters of 1 molar Tris pH 8.0, 50 microliters 8 molar Urea, and 50 microliters 150 millimolar NaCl 50 millimolar Tris pH 8.0 containing 100ng of Trypsin Gold (Promega). Tryptic digestion was allowed to proceed with nutation at 37 °C for between 6 and 18 hours before acidification with 10 microliters of 10% Formic Acid and 10 microliters of 10% Trifluoroacetic acid.

### Automated Affinity Purification

Cells were lysed the same way as for manual affinity purification, featuring the same sonification and centrifugation. Lysates were then transferred to 96-well plates with 2.2 milliliter square wells (NEST 503021). Additional plates were prepared containing ALFA beads (4 microliters of slurry into 100 microliters mRIPA lysis buffer, also containing a 96-tip comb, NEST 503311), two mRIPA lysis buffer washes (500 microliters buffer in each well), one water wash (one plate, 1000 microliters sterile water per well), and acid elution buffer (100 microliters per well). The plates were placed into a KingFisher MagMAX Express 96 instrument configured with BindIt 4.1 software to perform affinity purification for up to 96 samples at a time. The method file is included in the supplemental materials. Elutions were transferred from the 96-well plate to PCR tube strips using a multichannel pipette where they were subject to tryptic digestion with conditions identical to that of manual purification.

### MS Sample preparation

Acidified tryptic digests were desalted using 20mg of C18 resin extracted from 200mg SepPak cartridges (Waters) placed into Pierce Micro Spin columns (Thermo Scientific). Resin was conditioned with 150 microliters of 80% ACN 0.1% TFA and equilibrated with 150 microliters of 0.1% TFA before applying acidified tryptic digests. Resin was washed with 350 microliters of 0.1% acetic acid before peptides were eluted into a 1.5 milliliter microcentrifuge tube with 100 microliters of 80% ACN 0.1% acetic acid. All steps were performed using centrifugation at 800G for 60 seconds. Elutions were dried via vacuum concentrator and resuspended in 2 microliters of LC-grade water before autosampler injection for LC-MS/MS analysis.

### MS Data Acquisition

Data-dependent analyses for spectral library building were performed on a Q Exactive HF Orbitrap mass spectrometer using a 75-minute method containing a 40-minute reverse-phase gradient. This method featured a full MS scanning resolution of 60,000, an AGC target of 3e6, a maximum IT of 30ms, and a scan range of 380 to 1800 m/z. Data-dependent Top-20 MS2 parameters featured a resolution of 15,000, an AGC target of 1e5, a maximum IT of 100ms, an isolation window of 2.0 m/z, and an NCE of 28. Parallel reaction monitoring analyses for quantitative analyses were performed on the same instrument using a 33-minute method containing a 10-minute reverse-phase gradient. This method featured an MS2 resolution of 15,000, an AGC target of 2e5, a maximum IT of 120ms, an isolation window of 0.4 m/z, and an NCE of 28. Inclusion lists featured scheduling with 1.5-minute-wide retention time windows.

### Phosphoproteomics

Pellets were subject to nuclear enrichment via resuspension in 2 milliliters of hypotonic lysis buffer and incubated on ice for 20 minutes. Nuclei were pelleted via centrifugation for 5 minutes at 1000G. The nuclear pellets were then resuspended in 2 milliliters of mRIPA lysis buffer and lysed via sonification using a Branson Probe tip sonifier set to 15% amplitude with three 5-second pulses. Lysates were then transferred to 50 milliliter ultracentrifuge tubes for 30 minutes of centrifugation at 45,000G kept at 4□. Bradford assays were performed to measure protein concentration, after which lysates for each SILAC channel were pooled for a total volume of 3mL. Pooled lysates were then denatured and reduced with 1% SDS and 5 millimolar DTT at 42°C for 15 minutes, and then alkylated with 25 millimolar iodoacetamide at room temperature for 15 minutes. Lysates were mixed with 10 milliliters of cold PPT solution (49.9% EtOH, 50% acetone, 0.1% acetic acid) to precipitate on ice for 30 minutes, after which precipitated protein was pelleted via centrifugation at 4000G for 5 minutes. Pellets were resuspended with 1 milliliter of deionized water containing 10 microliters of Urea/Tris solution (8 molar urea, 50 millimolar Tris pH 8.0). Suspensions were transferred to clean ultracentrifuge tubes and centrifuged at 45,000G for 5 minutes. Supernatant was aspirated before 1 milliliter of deionized water was gently added to the ultracentrifuge tube and “rolled over” the pellet three times to remove residual contaminants. This water was aspirated and the pellet itself was resuspended in 2 milliliters of Urea/Tris solution. The resuspended pellet was transferred to a 15 milliliter centrifuge tube and 6 milliliters of NaCl/Tris solution was added (150 millimolar NaCl, 50 millimolar Tris pH 8.0). This resuspension was sonified for 10 seconds at 15% amplitude to break up pieces of precipitated protein. 40 microliters of 1 milligram per milliliter TPCK-treated trypsin was added, after which the tube was parafilmed and nutated at 37□ for 12 hours.

### Database Searching

Raw MS/MS spectra were searched using the Comet search engine (part of the Trans Proteomic Pipeline 7.0.0; Seattle Proteome Center) over a composite human protein database consisting of the *Homo sapiens* proteome and the sequence of the ProKAS module^48^. Search parameters allowed for semi-tryptic peptide ends, a mass accuracy of 15 ppm for precursor ions, variable modifications for SILAC lysine and arginine (8.0142 and 10.00827 daltons, respectively), variable modification for STY phosphorylation (79.966331 daltons), and a static mass modification of 57.021465 daltons for alkylated cysteine residues. Phosphorylation site localization probabilities were determined using PTMProphet, with SILAC quantification of identified phosphopeptides being performed using XPRESS (both modules part of the Trans Proteomic Pipeline; Seattle Proteome Center) ^49^. Phosphoproteomic data was subject to Bowtie filtering as previously described in Faca et al. 2020 ^50^. Processed phosphoproteomic data is available in supplemental table X. Library-building DDA data searched similarly, with search results being imported into Skyline (MacCoss Lab) for library curation.

### Skyline Data Analysis

PRM data files were converted to mzXML format via msConvert (Proteowizard) and imported into Skyline documents featuring parameters that are present in the Sky.zip file included in the supplement^51,52^. Transitions were checked manually for co-elution before exporting a report as a .csv containing Protein, Peptide Modified Sequence, RatioLightToHeavy, Normalized Area, and Replicate. Data in these reports were then processed such that SILAC ratios for each phosphopeptide were normalized against the SILAC ratios for their unphosphorylated counterparts to account for channel abundance differences. For label free experiments, peak areas for phosphorylated peptides were first divided by peak areas for unphosphorylated peptides. The quotients for treated samples were then compared to that of untreated samples to acquire quantitative ratios. Error bars and envelopes reflect the standard deviation of three biological replicates unless otherwise noted.

### Fluorescence microscopy

U2OS cells (ATCC) were transfected and incubated for 24 hours before treatment with 10 micromolar 5-Ethynyl-2′-deoxyuridine (EdU; Sigma, 900584) for 15 minutes. Cells were washed once with ice-cold phosphate-buffered saline (PBS) and fixed with 4% paraformaldehyde (PFA; Sigma, 158127) for 10 minutes at room temperature. Following fixation, cells were washed twice with PBS, permeabilized with 0.5% Triton X-100 in PBS (Sigma, T9284) for 10 minutes, and subsequently washed twice with 3% bovine serum albumin (BSA; Roche Diagnostics, 03117332001) in PBS. EdU detection was performed using click chemistry, where cells were incubated with a reaction mixture containing10 millimolar sodium-L-ascorbate (Sigma, A7631), 0.1 millimolar Atto 594 azide (Sigma, 72998), and 5 millimolar copper (II) sulfate (Sigma, C1297) for 30 minutes. The reaction solution was removed, and cells were washed three times with PBS, followed by incubation with 5% BSA for 30 minutes at room temperature. Next, cells were incubated with anti-GFP antibody (1:200; Santa Cruz, sc-9996) overnight at 4 □, followed by three washes with PBS coverslips were mounted using Vectashield with DAPI (Vector Laboratories, H-2000) and sealed with nail polish. Imaging was performed on a Leica DMI8 microscope equipped with an HC PL APO 100x/1.40 OIL objective. Image acquisition and processing were conducted using Leica Application Suite X (LASX) software (ver. 3.8.2.27713). Colocalization analysis between ProKAS-PCNA and EdU signals was performed using Fiji/ImageJ (NIH). A randomly drawn line across the nucleus was used to generate intensity profiles via the Plot Profile function in Fiji/ImageJ. Data were then imported into GraphPad Prism 9 for visualization.

### *De novo* Design of CHK1 Kinase-Specific Sensor

To identify optimal CHK1 sensor peptide sequences, we employed a genetic algorithm leveraging PSPA-based kinase preference scores. The algorithm aimed to maximize the predicted phosphorylation of a given 10-amino acid sequence by CHK1 while minimizing its preference for other kinases in the PSPA dataset^20^. We initialized the algorithm with a population of 50-200 randomly generated sequences and allowed it to evolve over 10-50 generations. In each generation, the fitness of each sequence was evaluated based on its PSPA scores for all 303 kinases in the dataset. The fittest sequences were then selected for reproduction through crossover and mutation operations, creating a new population for the next generation. This iterative process led to the convergence of the population towards sequences with high predicted CHK1 specificity. The algorithm was run multiple times, varying the start population size and the number of generations, to explore a wide sequence space. The resulting list of optimized peptides was further curated to ensure amino acid diversity among the final 10 selected candidates. Each sequence was flanked by AGA and R residues to ensure efficient trypsin digestion and generation of unique detectable tryptic peptides. A synthetic gBlock encoding the 10 candidate peptide sequences was obtained from IDT and cloned into a ProKAS vector containing the 53BP1 nuclear localization signal (NLS) to target the expressed biosensor to the nucleus.

For experimental validation, HEK293T cells were transfected with the ProKAS vector containing the multiplexed CHK1 sensor candidates. Following transfection, cells were treated with genotoxic agents (e.g., camptothecin or hydroxyurea) to induce CHK1 activation. Cell lysates were collected, and the ProKAS sensors were immunoprecipitated using anti-ALFA beads. The immunoprecipitants were then trypsin digested, and the resulting peptides were analyzed by mass spectrometry. The phosphorylation levels of each candidate peptide were quantified, and the peptide demonstrating the highest inducibility and specificity upon CHK1 activation was selected as the final CHK1 sensor.

### Immunoblotting

Cells were harvested and lysed in mRIPA lysis buffer as described above. Lysates were cleared by centrifuging at 13,000G for 5 minutes and then mixed with 3X SDS sample buffer (90 millimolar Tris-HCl pH 7.5, 3% SDS, 30% glycerol, 0.03% bromophenol blue, 60 millimolar DTT). 10% and 14% SDS-PAGE gels were run at 30 milliamps per gel before performing a wet transfer to polyvinyulidene difluoride (PVDF) membranes. Membranes were incubated with primary antibodies overnight at 4□ before being developed using BioRad Clarity ECL substrate and ChemiDoc Imaging system. The following antibodies were used: anti-KAP1 Bethyl A700-014-T, anti-pKAP1-S824 Bethyl A304146AT, anti-CHK1 Santa Cruz Biotechnology SC-8408, anti-pCHK1-S345 Cell Signaling 2341S, anti-Beta-Actin Proteintech 66009-1-Ig.

## Supplemental Figures

**Figure S1:**
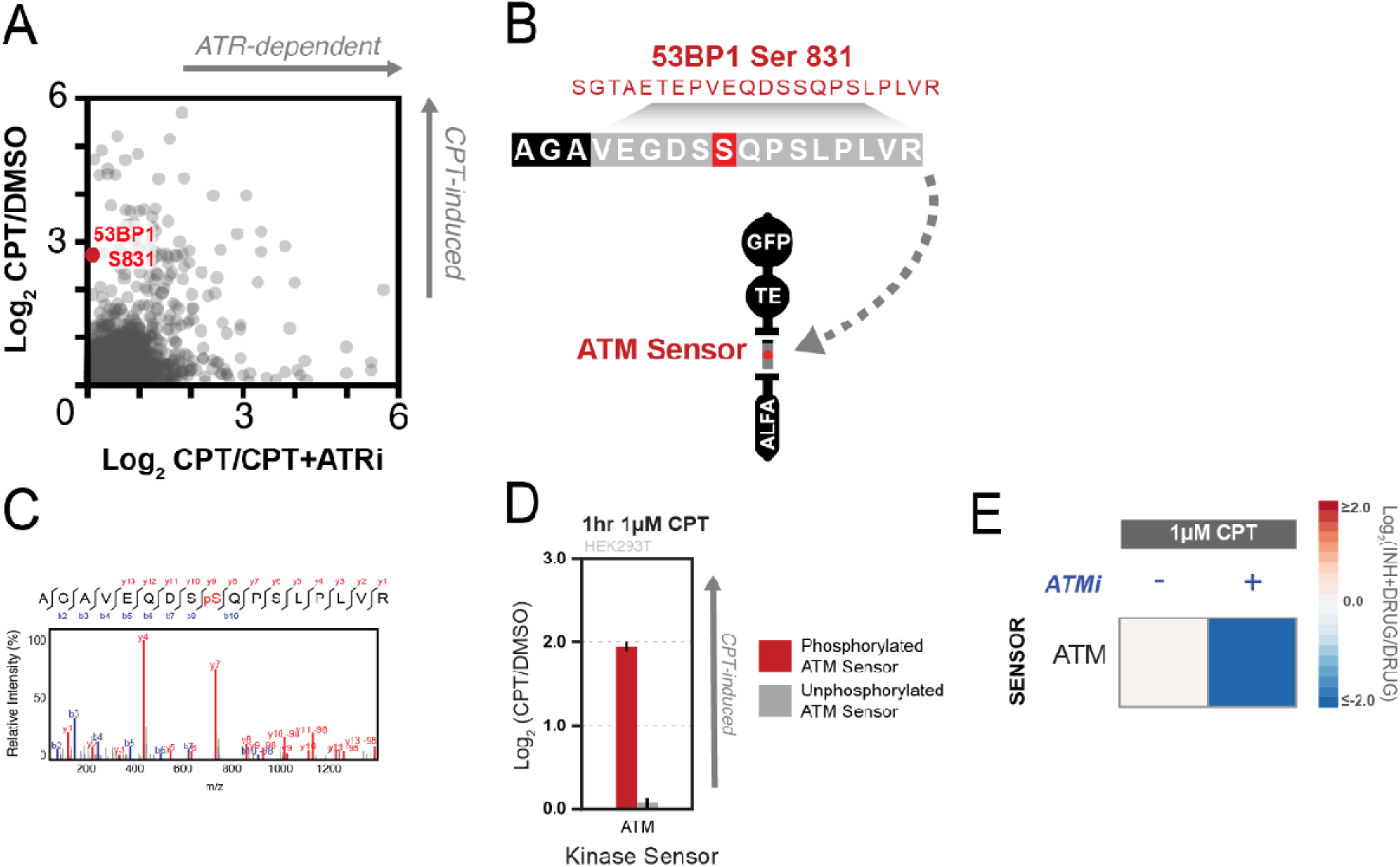
Process for ATM sensor curation. A) Phosphorylation site Serine 831 on 53BP1 was selected from the phosphoproteomic data generated to determine an ATR sensor. This site was induced by CPT but not dependent on ATR, and had previously been found to be phosphorylated by ATM. B) The 53BP1 Serine 831 phosphopeptide sequence was cloned into a ProKAS biosensor after adding a code to the N-terminal end. C) Mass spectrometry was able to detect the phosphorylated form of this ATM sensor candidate, as shown by the MS/MS spectrum. D) Phosphorylation of this ATM sensor candidate was observed to increase after cells were treated with 1 micromolar CPT for 1 hour, whereas the unphosphorylated form of this sensor did not change in abundance. E) Induced phosphorylation of this ATM sensor candidate did not occur after the addition of the ATM inhibitor AZD-0156. Error bars/envelopes in D indicate the mean and standard deviation of triplicate independent experiments.

**Figure S2:**
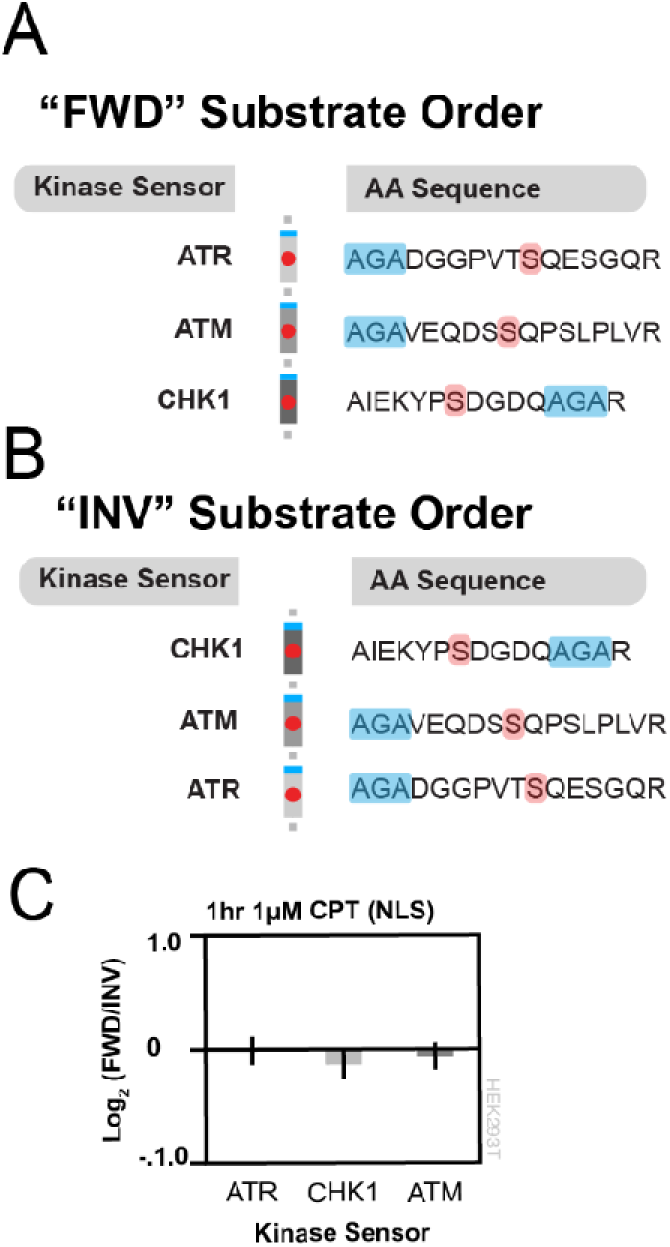
ProKAS biosensor designs for testing the effect of kinase sensor order on phosphorylation levels. A) The “forward” order of the kinase sensors: ATR - ATM - CHK1. B) The “inverted” order of the kinase sensors: CHK1 - ATM - ATR. C) Forward and inverted ProKAS biosensors showed no difference in sensor phosphorylation after 1 hour of treatment with 1 micromolar CPT. Error bars/envelopes in C indicate the mean and standard deviation of triplicate independent experiments.

**Figure S3:**
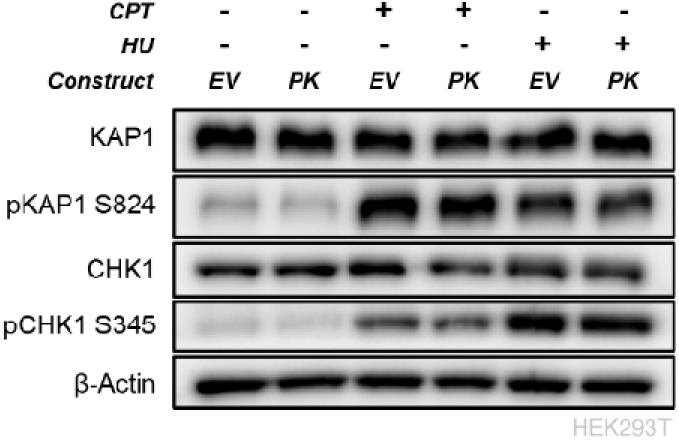
Kinase sensor expression does not impact endogenous DDR signaling. Western blot where markers for ATM (KAP1 phosphorylation at S824) and ATR (CHK1 phosphorylation at S345) were measured in cells expressing an “EV” version of the ProKAS biosensor or the version containing kinase sensors for ATR, ATM, and CHK1 (denoted as “PK”). “EV” biosensors include all components of the ProKAS biosensor except for the MKS. Cells were treated with 1 micromolar CPT or 1 millimolar HU for 2 hours before lysis and Western blotting.

**Figure S4:**
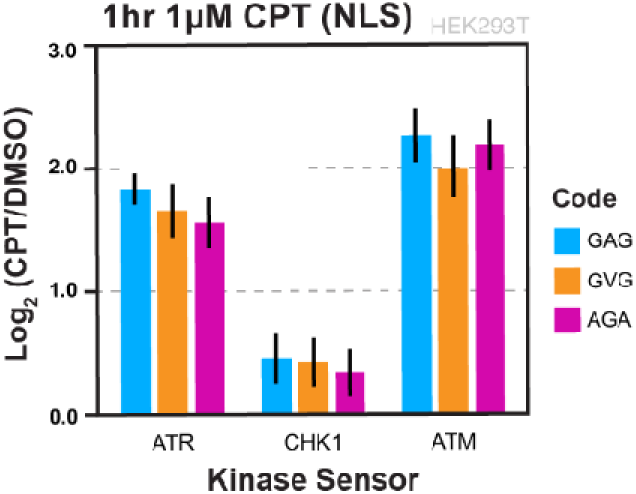
Amino acid barcodes show no difference in phosphorylation efficiency. Nuclear ProKAS constructs containing sensors for ATR, CHK1, and ATM with three different amino acid barcodes (GAG, GVG, and AGA) were co-expressed in HEK293T cells and showed no significant difference in levels of induced phosphorylation after treatment with 1 micromolar CPT for 1 hour. Error bars/envelopes indicate the mean and standard deviation of triplicate independent experiments.

**Figure S5:**
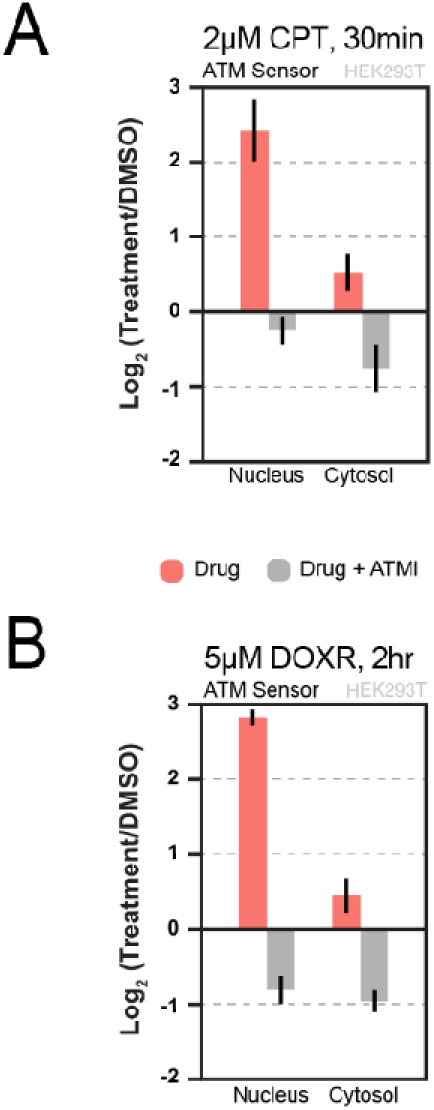
Inhibition of nuclear and cytosolic ATM activity. A) ATM inhibitor impairs ATM sensor phosphorylation in both the nucleus and cytosol after treatment with CPT (Label free quantification). B) ATM inhibitor impairs ATM sensor phosphorylation in both the nucleus and cytosol after treatment with DOXR (Label free quantification). Error bars/envelopes indicate the mean and standard deviation of duplicate independent experiments for A and triplicate independent experiments for B.

