## Supplementary material for "Proteomic Sensors for Quantitative, Multiplexed and Spatial Monitoring of Kinase Signaling": de novo algorithm script: GA-final-new-PSSM_SCORING-MN.py

```
# Import necessary libraries
import random  # For generating random numbers
import numpy as np  # For numerical operations
import pandas as pd  # For data manipulation and analysis
import os  # For interacting with the operating system
import matplotlib.pyplot as plt  # For creating 
from deap import base, creator, tools, algorithms  # For Genetic Algorithm implementation
import csv  # For working with CSV files
import time  # For measuring time
import glob  # For finding files that match a pattern

# ---------------------------- Parameters, Inputs, outputs ----------------------------#
dir_path = os.path.dirname(os.path.realpath(__file__))

# Set paths to directories and files
main_directory = os.chdir("Users/will/Desktop/Smolka Lab/SELiKS Project/ProKAS Algorithm")
pssm_directory = "Users/will/Desktop/Smolka Lab/SELiKS Project/ProKAS Algorithm" # Directory containing PSSM files
output_file_directory = "Users/will/Desktop/Smolka Lab/SELiKS Project/ProKAS Algorithm"  # Directory to save output files
output_temp_name = output_file_directory + "temp.csv"  # Temporary output file

# Dictionary to store positions excluded from certain amino acids
excluded_positions = {
    0: [],
    1: [],
    2: [],
    3: [], 
    4: [],
    6: [],
    7: [],
    8: [],
    9: []
}

# Target kinase for optimization
target_kinase_filename = "DNAPK"

# List of CKS kinases
CKS_kinases = ['ATR', 'ATM', 'CHK1', 'CHK2']

# Peptide Score Calculation 
Weight_CKS_kinases = 1
Weight_Background_kinases = 9

# Genetic Algorithm parameters
pop_size = 20  # Population size
n_gen = 10  # Number of generations
cycles = 10  # Number of optimization cycles

# ------------------------------------ Functions ------------------------------------#

# Function to load PSSM matrices from CSV files
def load_pssms(directory):
    pssms = {}  # Dictionary to store PSSMs
    for filename in os.listdir(directory):  # Iterate through files in the directory
        if filename.endswith('.csv'):  # Check if file is a CSV file
            kinase_name = filename[:-4]  # Extract kinase name from filename
            pssm = pd.read_csv(os.path.join(directory, filename), index_col=0)  # Load PSSM into DataFrame
            pssm.columns = range(-5, 5)  # Set column names to represent positions
            pssms[kinase_name] = pssm  # Store PSSM in the dictionary
    return pssms

# Function to calculate maximum and minimum possible scores for each kinase
def calculate_max_min_scores(pssms):
    max_scores = {}
    min_scores = {}
    for kinase, pssm in pssms.items():
        max_sum = np.sum([pssm[col].max() for col in pssm.columns])  # Sum of maximum values in each column
        min_sum = np.sum([pssm[col].min() for col in pssm.columns])  # Sum of minimum values in each column
        max_scores[kinase] = max_sum
        min_scores[kinase] = min_sum
    return max_scores, min_scores

# Function to create an individual (peptide sequence) for the GA
def create_individual():
    individual = [random.choice(positions[i]) for i in range(len(positions))]  # Randomly select amino acids for each position
    individual[5] = 'S'  # Fix the 6th position to 'S'
    return individual

# Function to score a peptide based on PSSMs
def score_peptide(peptide, pssms, max_scores, min_scores):
    normalized_scores = []
    for pssm_name, pssm in pssms.items():
        raw_score = np.sum([pssm.at[amino_acid, pos - 5] for pos, amino_acid in enumerate(peptide)])  # Calculate raw score
        normalized_score = (raw_score - min_scores[pssm_name]) / (max_scores[pssm_name] - min_scores[pssm_name])  # Normalize score
        normalized_scores.append(normalized_score)
    return normalized_scores

# Function to evaluate the fitness of an individual
def evaluate(individual, pssms, target_kinase_filename, max_scores, min_scores):
    peptide = ''.join(individual)  # Join amino acids into a peptide sequence
    scores = score_peptide(peptide, pssms, max_scores, min_scores)  # Get scores for all kinases
    target_kinase_index = list(pssms.keys()).index(target_kinase_filename)  # Find index of the target kinase
    non_target_CKS_scores = [scores[list(pssms.keys()).index(item)] for item in CKS_kinases if item != target_kinase_filename]  # Scores for non-target CKS kinases
    target_score = scores[target_kinase_index]  # Score for the target kinase
    CKS_score = np.mean(non_target_CKS_scores)  # Average score for non-target CKS kinases
    CKS_std = np.std(non_target_CKS_scores)  # Standard deviation of non-target CKS kinase scores
    Background_scores = np.mean([score for i, score in enumerate(scores) if i != target_kinase_index])  # Average score for background kinases
    Background_std = np.std([score for i, score in enumerate(scores) if i != target_kinase_index])  # Standard deviation of background kinase scores

    # Calculate fitness components
    Target_to_Background_score = (target_score - Background_scores) / Background_std
    Target_to_CKS_score = (target_score - CKS_score) / CKS_std

    # Weights for fitness components
    Weight_CKS = Weight_CKS_kinases
    Weight_Background = Weight_Background_kinases

    # Calculate final fitness
    Fitness = (Weight_CKS * Target_to_CKS_score) + (Weight_Background * Target_to_Background_score)
    return (Fitness,)  # Return fitness as a tuple

# Function to run the Genetic Algorithm
def run_ga(pop_size, n_gen):
    population = toolbox.population(n=pop_size)  # Create initial population
    fitness_history = []  # List to store fitness history

    for gen in range(n_gen):  # Iterate through generations
        offspring = algorithms.varAnd(population, toolbox, cxpb=0.8, mutpb=0.8)  # Create offspring using variation operators
        invalid_ind = [ind for ind in offspring if not ind.fitness.valid]  # Find individuals without fitness values
        fitnesses = map(toolbox.evaluate, invalid_ind)  # Evaluate fitness for invalid individuals
        for ind, fit in zip(invalid_ind, fitnesses):
            ind.fitness.values = fit  # Assign fitness values

        population = toolbox.select(offspring + population, pop_size)  # Select individuals for the next generation

        # Handle cases where fitness values are missing or invalid
        for ind in population:
            if not isinstance(ind.fitness.values, tuple) or len(ind.fitness.values) == 0:
                ind.fitness.values = (float('-inf'),)

        # Calculate and store fitness statistics
        fitness_values = [ind.fitness.values[0] for ind in population]
        avg_fitness = np.mean(fitness_values)
        best_fitness = max(fitness_values)
        fitness_history.append((avg_fitness, best_fitness))

        # Plot fitness history
        plt.figure(figsize=(10, 5))
        plt.plot([f[0] for f in fitness_history], label=f"Avg Fitness (gen {gen}): {avg_fitness:.2f}")
        plt.xlabel("Generation")
        plt.ylabel("Fitness")
        plt.title("Fitness Over Generations")
        plt.legend()
        plt.show()

    return population, fitness_history, avg_fitness

# Function to filter peptides based on excluded positions and output to CSV
def filter_and_output_peptides(population, excluded_positions, file_name):
    # Function to check if a peptide is valid based on excluded positions
    def is_peptide_valid(peptide, excluded_positions):
        for pos, excluded_aas in excluded_positions.items():
            if peptide[pos] in excluded_aas:
                return False
        return True

    valid_population = [peptide for peptide in population if is_peptide_valid(peptide, excluded_positions)]  # Filter valid peptides

    # Write valid peptides and their scores to CSV
    with open(file_name, 'w', newline='') as csvfile:
        fieldnames = ['Peptide Rank', 'Sequence'] + list(pssms.keys())
        writer = csv.DictWriter(csvfile, fieldnames=fieldnames)
        writer.writeheader()
        for rank, peptide in enumerate(valid_population, start=1):
            peptide_with_S = peptide[:5] + ['S'] + peptide[6:]  # Insert 'S' at the 6th position
            peptide_scores = score_peptide(''.join(peptide_with_S), pssms, max_scores, min_scores)  # Score the peptide
            row = {'Peptide Rank': f'Peptide {rank}', 'Sequence': ''.join(peptide_with_S)}  # Create a row for the CSV
            row.update({kinase: score for kinase, score in zip(pssms.keys(), peptide_scores)})  # Add scores for each kinase
            writer.writerow(row)  # Write the row to the CSV

# Function to merge and process multiple CSV files
def merge_and_process_csv_files(base_name, output_file):
    file_pattern = f"{base_name}*.csv"  # Pattern to match CSV files
    files = glob.glob(file_pattern)  # Find files matching the pattern
    files.sort()  # Sort files by name
    all_dataframes = []  # List to store DataFrames

    for i, file_name in enumerate(files):
        if os.path.exists(file_name):  # Check if file exists
            df = pd.read_csv(file_name, skiprows=0)  # Read CSV into DataFrame
            all_dataframes.append(df)  # Add DataFrame to the list
        else:
            print(f"File not found: {file_name}")  # Print error message if file is not found

    merged_dataframe = pd.concat(all_dataframes, ignore_index=True)  # Merge all DataFrames

    merged_dataframe.sort_values(by=target_kinase_filename, ascending=False, inplace=True)  # Sort by target kinase score in descending order

    merged_dataframe.to_csv(output_file, index=False)  # Save merged DataFrame to CSV
    merged_dataframe.drop('Peptide Rank', axis=1, inplace=True)  # Drop 'Peptide Rank' column

# ---------------------------- GA setup ----------------------------#

# Load PSSMs and calculate max/min scores
pssms = load_pssms(pssm_directory)
max_scores, min_scores = calculate_max_min_scores(pssms)

# Set up DEAP framework for Genetic Algorithm
if not hasattr(creator, "Fitness"):
    creator.create("Fitness", base.Fitness, weights=(1.0,))  # Create Fitness class
if not hasattr(creator, "Individual"):
    creator.create("Individual", list, fitness=creator.Fitness)  # Create Individual class
toolbox = base.Toolbox()
toolbox.register("individual", tools.initIterate, creator.Individual, create_individual)  # Register individual creation function
toolbox.register("population", tools.initRepeat, list, toolbox.individual)  # Register population creation function
toolbox.register("evaluate", evaluate, pssms=pssms, target_kinase_filename=target_kinase_filename, max_scores=max_scores, min_scores=min_scores)  # Register evaluation function
toolbox.register("mate", tools.cxUniform, indpb=0.1)  # Register crossover operator
toolbox.register("mutate", tools.mutShuffleIndexes, indpb=0.05)  # Register mutation operator
toolbox.register("select", tools.selTournament, tournsize=2)  # Register selection operator

# ---------------------------- Main ----------------------------#

if __name__ == "__main__":
    # Define possible amino acids for each position (with 'S' fixed at position 5)
    positions = ['MAGVLIFWYDEQNRKSTPH', 'MAGVLIFWYDEQNRKSTPH', 'MAGVLIFWYDEQNRKSTPH', 'MAGVLIFWYDEQNRKSTPH', 'MAGVLIFWYDEQNRKSTPH', 
                 'S', 'MAGVLIFWYDEQNRKSTPH', 'MAGVLIFWYDEQNRKSTPH', 'MAGVLIFWYDEQNRKSTPH', 'MAGVLIFWYDEQNRKSTPH']

    count = 0
    while count < cycles:  # Run multiple optimization cycles
        output_file_name = output_file_directory + f"{target_kinase_filename}-top_peptides_{count+1}.csv"  # Output file name
        start_time = time.time()  # Record start time
        final_population, fitness_history, avg_fit = run_ga(pop_size, n_gen)  # Run GA
        best_individual = tools.selBest(final_population, 1)[0]  # Get the best individual
        duration = time.time() - start_time  # Calculate duration
        print(f"Process {count+1} took {duration:.2f} secs (Final Avg fitiness: {avg_fit:.2f})")

        # Filter and output peptides to temporary file
        filter_and_output_peptides(final_population, excluded_positions, output_temp_name)

        # Read, clean, and reorder columns in the temporary file
        df = pd.read_csv(output_temp_name)
        df_cleaned = df.drop_duplicates(subset='Sequence')
        fixed_cols = ['Peptide Rank', 'Sequence']
        cols = fixed_cols + CKS_kinases + [col for col in df_cleaned.columns if col not in CKS_kinases + fixed_cols]
        df_reordered = df_cleaned[cols]
        df_reordered.to_csv(output_file_name, index=False)  # Save to final output file

        count = count + 1
        total_duration = 0 + duration

    # Merge and process all output files into a final file
    base_name_pattern = output_file_directory + target_kinase_filename + '-top_peptides_'
    output_file_name = output_file_directory + target_kinase_filename + '_final.csv'
    merge_and_process_csv_files(base_name_pattern, output_file_name)

    print(f"Total execution time: {total_duration:.2f} minutes")
                                                         
```
